# Early removal of the infrapatellar fat pad beneficially alters the pathogenesis of moderate stage idiopathic knee osteoarthritis in the Dunkin Hartley guinea pig

**DOI:** 10.1101/2022.02.04.479182

**Authors:** Maryam F. Afzali, Lauren B. Radakovich, Margaret A. Campbell, Joseph L. Sanford, Madeline M. Sykes, Angela J. Marolf, Tammy H. Donahue, Kelly S. Santangelo

## Abstract

**Background:** The infrapatellar fat pad (IFP) is the largest adipose deposit in the knee; however, its contributions to the homeostasis of this organ remain unknown. To determine the influence of this depot on joint health, this study determined the progression of osteoarthritis (OA) following excision of the IFP in a rodent model of naturally-occurring disease.

**Methods:** Male Dunkin-Hartley guinea pigs (n=10) received surgical removal of the IFP in one knee at 3 months of age; contralateral knees received sham surgery as matched internal controls. Treadmill-based gait analysis was performed prior to IFP removal and monthly thereafter. Animals were harvested at 7 months of age. Both knees were processed for microcomputed tomography (microCT), histopathology, transcript expression analyses, and immunohistochemistry (IHC).

**Results:** Fibrous connective tissue (FCT) developed in place of the native fat pad. Gait demonstrated no significant differences between IFP removal and contralateral limbs. MicroCT OA scores were improved in knees with the FCT. Histopathology confirmed maintenance of articular cartilage structure, proteoglycan content, and chondrocyte cellularity in FCT-containing knees. Transcript analyses revealed decreased expression of adipose-related molecules and inflammatory mediators in FCTs compared to IFPs. This was corroborated via IHC for select inflammatory mediators.

**Discussion/Conclusion:** Formation of the FCT resulted in reduced OA-associated changes in both bone and cartilage. A decrease in inflammatory mediators at transcript and protein levels may be associated with these improvements. The IFP may therefore play a role in the pathogenesis of knee OA in this strain, with removal prior to disease onset appearing to have short-term benefits.

## Introduction/Background

Primary osteoarthritis (OA), particularly knee OA, currently burdens greater than 273 million people globally^1^. As a leading cause of pain and disability, OA is a major contributor to decreased quality of life^2^. Consequently, more than 1 million people undergo knee arthroscopy or joint replacement surgery each year due to end-stage OA in the United States^3^, alone, with the annual economic loss to Americans approaching $200 billion^4^. Unfortunately, there are no therapeutic regimens able to restore damaged cartilage to its normal phenotype or slow the progression of joint degeneration^5^. This reflects a need to improve our current understanding of pathophysiology of the disease, which is associated with both inflammatory and biomechanical causes^6^.

The knee is composed of many tissue types, including a number of continuous but distinct adipose depots; specifically, the infrapatellar fat pad (IFP), the posterior knee fat pad, the quadriceps fat pad, and the pre-femoral fat pad^7^. The IFP is the largest of these and is found in the anterior aspect of the joint in the space shaped by the patella, femoral condyles, and tibial plateau. In spite of its bulk, the exact functions of the IFP are not completely understood. The main role is thought to be facilitating the distribution of synovial fluid across the knee joint, thereby providing lubrication^8–10^. It also likely supplies shock absorbance from mechanical forces, knee joint stability, and may prevent instability and/or injury associated with loading forces to the knee joint^10,11^. However, *ex vivo* work performed on cadavers revealed that resection of the IFP decreased tibial rotation of the knee^12^. From this, the authors extrapolated that the IFP may contribute to dictating the range of motion of the knee joint.

Despite this uncertainty, the IFP is considered a player in overall knee joint homeostasis and there is evidence to support its part in the pathogenesis of knee joint OA^8–15^. In particular, the IFP is comprised of a network of adipocytes, fibroblasts, leukocytes (primarily macrophages and lymphocytes), and collagen matrix^7–9^. As such, it is positioned to be a source of inflammatory mediators and/or immune modulation that may contribute to OA^7–16^. For example, a study utilizing human tissues demonstrated that IFP-derived adipocyte conditioned media induced a pro-inflammatory response in T lymphocytes that resulted in increased proliferation and cytokine production^16^. In addition, Distal et al. have shown that interleukin (IL)-6 secretion from the IFP of women with knee OA was more than twice that of subcutaneous thigh fat^17^.

Hoffa’s disease, sometimes called hoffitis, is a disease of the IFP (also been referred to as Hoffa’s fat pad). The pathophysiology of this disorder is not well documented; however, several mechanisms have been implicated, including acute trauma, microtrauma, and over-solicitation (repeated rotation and hyperextension)^18^. Regardless of the inciting cause, patients present with limitation of knee extension. Subsequent magnetic resonance imaging highlights signal abnormalities in the IFP along the path of the adipose ligament^19–21^. Conservative approaches are proposed as a first-line of therapy; however, if response to these options is unsuccessful and/or the disease becomes chronic, arthroscopic resection/removal remains the next treatment of choice. In particular, individuals with this condition experience pain relief from undergoing arthroscopic subtotal removal of this adipose depot^22^.

Given the above, we aimed to determine if unilateral removal of the IFP would alter the pathogenesis of knee joint OA in an animal model of primary disease. The aims of this study were two-fold: 1) to determine whether gait changes, an indication of possible symptom modification, would occur; and 2) to assess potential disease modification. We employed the Hartley guinea pig for this study, as it is a valuable rodent model of idiopathic OA with histopathologic lesions similar to those seen in people^23,24^. We postulated that the IFP acts as a local source of inflammation in this model^23^ and that, by surgically removing it, we might improve OA outcomes by eliminating a source of negative mediators.

## Methods

### Animals

All procedures were approved by the Institutional Animal Care and Use Committee and performed in accordance with the NIH Guide for the Care of Use of Laboratory Animals. Sample size was determined from a pilot study focused on histologic assessment of OA as the primary outcome. Using a within group error of 0.5 and a detectable contrast of 0.5 in a linear regression model, power associated with an alpha level of 0.05 was calculated as 0.95 with a sample size of 10. Thus, 10 8-week-old male Dunkin-Hartley guinea pigs were purchased from a commercial vendor (Charles River Laboratories, Wilmington, MA). Animals were maintained at the University’s Laboratory Animal Resources housing facilities and were monitored daily by a veterinarian. All guinea pigs were singly-housed in solid bottom cages and provided standard chow, hay cubes, and water *ad libitum*. Sixteen male Hartley guinea pigs of the same age, from a coincident but unrelated project, were utilized as an untreated control group for body weight comparisons.

### Surgical removal of the IFP

Resection of the IFP was performed on 12-week-old animals. After medial parapatellar arthrotomy of the right knee, the patella was displaced cranially with the knee in extension to permit access to the femoral groove. The IFP was then exposed medially to allow dissection and removal. The patella was repositioned and the skin incision closed. An identical sham procedure, with minor manipulation but without removal of the IFP, was performed on the left knee and served as a matched internal control for each animal.

### Treadmill-based gait analysis

Obligatory gait analysis was performed using a DigiGait™ treadmill system (Mouse Specifics, Inc., Framingham, MA, USA). Animals were acclimated to the apparatus over a 2-week period. Data collection was performed by the same handlers during the same time period (8:00AM to 12:00PM); the order in which animals were analyzed was randomly selected. Baseline gait analysis was performed the day before IFP removal surgery. Subsequent data were collected every 4 weeks after surgery, with the final time point occurring the day before termination.

### Tissue collection

Animals were harvested at 7 months of age. Body weights were recorded; animals were transferred to a CO_2_ chamber for euthanasia. Hind limbs were removed at the coxofemoral joint. The lengths of the left and right femurs were measured using calipers. The left and right hind limbs were then placed in 10% neutral buffered formalin for 48 hours and transferred to phosphate buffer saline (PBS) for microcomputed tomography (microCT) analysis. After microCT, limbs were placed in 12.5% solution of ethylenediaminetetraacetic acid (EDTA) at pH 7 for decalcification. EDTA was replaced twice weekly for 6 weeks.

### MicroCT

Knee joints were scanned in PBS using the Inveon microPET/CT system (Siemens Medical Solutions, Malvern PA), with a voxel size of 18 μm, a voltage of 100 kV, and an exposure time of 1356 ms. Clinical features of boney changes of OA were scored using a published whole joint grading scheme^25^. Features graded include: presence/size and location of osteophytes, subchondral bone cystic changes, subchondral bone sclerosis, articular bone lysis, and intraarticular soft tissue mineralization. Images were scored by a board-certified veterinary radiologist (AJM) blinded to limb identification.

### Histologic Grading of OA

Following decalcification and paraffin-embedding, three sagittal 5μm sections were made through each knee joint: (i) mid-sagittal slices were used for histologic evaluation of the IFP; and sagittal slices through both the (ii) medial and (iii) lateral compartments were utilized to assess OA changes in four sites (medial tibia, lateral tibia, medial femur, and lateral femur) using OARSI recommended criteria^24^. The mid-sagittal sections were stained with hematoxylin and eosin (H&E) and Masson’s trichrome stain to confirm structural modifications. The medial and lateral compartments were stained with toluidine blue for OA grading. The OARSI published grading scheme is based upon species-specific features of OA including: articular cartilage structure, proteoglycan content, cellularity, and tidemark integrity. Two blinded veterinary pathologists (LBR and KSS) performed histological grading. Scores from each of the four anatomic locations were summed to obtain a total knee joint OA score for each right and left hind limb. An intraclass correlation coefficient of 0.9 for between-reviewer consistency was calculated; the few minor discrepancies identified were resolved prior to statistical analysis.

### Gene Expression using Nanostring technology

Total ribonucleic acid (RNA) was isolated from either the IFP or replacement tissue that remained in formalin-fixed paraffin-embedded blocks following acquisition of adequate sections for histopathology and immunohistochemistry (IHC) using a commercially available kit specifically designed for such (Roche, Basel, Switzerland). A custom set of guinea pig-specific probes were designed and manufactured by NanoString Technologies (Seattle, WA) for the following genes: adiponectin (ADIPOQ), leptin (LEP), and fatty acid synthetase (FASN), nuclear factor kappa-B transcription complex (NF-kB p65 & NF-kB p50), nuclear receptor subfamily 4 group A member 2 (NR4A2), monocyte chemoattractant protein-1 (MCP-1), complement component 3 (C3), and matrix metallopeptidase-2 (MMP-2) (Supplemental Table 1). Per initial RNA quantification (Invitrogen Qubit 2.0 Fluorometer and RNA High Sensitivity Assay Kit, Thermo Fisher Scientific, Waltham, MA) and Fragment Analyzer quality control subsets (Fragment Analyzer Automated CE System and High Sensitivity RNA Assay Kit, Agilent Technologies), the optimal amount of total RNA (800.00ng) was hybridized with the custom code-set in an overnight incubation set to 65°C, followed by processing on the NanoString nCounter FLEX Analysis system. Results were reported as absolute transcript counts normalized to positive controls and three housekeeping genes (b-actin, succinate dehydrogenase, and glyceraldehyde-3-phosphate dehydrogenase). Any potential sample input variance was corrected by use of the housekeeping genes and application of a sample-specific correction factor to all target probes. Data analysis was conducted using nSolver software (NanoString Technologies).

### IHC and quantitative analysis

IHC was performed on mid-sagittal sections containing the IFP or replacement tissue using polyclonal rabbit antibodies to MCP-1 (Abcam ab9669) or NF-κB p65 (Abcam ab86299), each at a concentration of 2.5 μg/ml. Prior to staining, slides were incubated in citrate buffer for 5 hours at 55°C for antigen retrieval, as recommend for skeletal tissues^26^. 2.5% normal goat serum was used as a blocking reagent. Slides were incubated in primary antibody overnight in a humidified chamber at 4°C, followed by a 30-minute incubation with a biotinylated goat anti-rabbit secondary antibody. Bone marrow hematopoietic cells and macrophages served as internal positive controls for each section. Negative assay controls – rabbit immunoglobulin at 2.5 μg/ml and secondary antibody, alone – did not result in background immunostaining. Sections were counterstained with hematoxylin, cover slipped, and imaged by light microscopy. Data was quantified as integrated optical density using ImagePro-Plus 7 Software (Media Cybernetics, Rockville, MD). Four 1-mm-wide regions of interest of the IFP and replacement tissue were analyzed for MCP-1 and Nf-kBp65 expression; data for each tissue type was averaged prior to statistical analysis.

### Statistical analyses

Rationale for excluding individual values from data sets were determined prior to analysis and included whether an appropriate sample was unable to be collected, did not pass quality control parameters, or if integrity was compromised. Four guinea pigs were only amendable for treadmill data collection on some, but not all, collection dates. Two animals did not have appropriate sections available for the whole joint OA score. One animal did not have adequate tissue available for either RNA extraction or IHC. Exclusion of these animals resulted in: n=6 animals for treadmill-based gait results; n=8 for OARSI scoring; and n=9 for transcript expression and IHC results. All remaining outcomes included n=10 animals.

Statistical analyses were performed with GraphPad Prism 9.1.1(La Jolla, CA) with significance set at p ≤ 0.05. Data underwent normality and variance testing with the Shapiro-Wilk test. Normally distributed data with similar variance were compared using paired t-test^†^ for normally distributed data; Wilcoxon matched-pairs signed rank test^◊^ was used for non-normally distributed data.

## Results

### General Description of Study Animals

All guinea pigs remained clinically healthy throughout the study. There was no significant difference in body weight when this cohort was compared to 7-month-old Hartley guineas pigs utilized as untreated controls in a parallel but unrelated study (Supplemental Figure S1A). Mean total body weight was 1098 g (95% CI: 1042-1155 g) in the IFP removal group and 1100 g (95% CI: 1070-1130 g) in the control group. To ensure that potential differences in gait were not attributable to changes in skeletal properties, left and right femurs lengths from all IFP removal guinea pigs were measured. Femur lengths between the left [sham control (mean=45.38 mm; 95% CI: 44.89-45.87 mm)] and right (IFP removal [mean= 45.37; 95% CI: 44.88-45.87 mm)] hind limbs were not significantly different (*P* = 0.2345) (Supplemental Figure S1B).

### Treadmill-based gait analyses

To assess whether removal of the IFP impacted the gait of each animal, we contrasted parameters for each matched hindlimb. The ventral view of a guinea pig (Figure 1A), digital video images (Figure 1B), and representative dynamic gait signals (Figure 1C) of a guinea pig walking on a transparent treadmill belt at 35 cm/s are provided. No differences in midline distance (Figure 1D), stride length (Figure 1E), swing (Figure 1F), stance (Figure 1G), brake (Figure 1H) or propel (Figure 1I) were seen between the IFP removal and contralateral hindlimbs.

**Figure 1.**
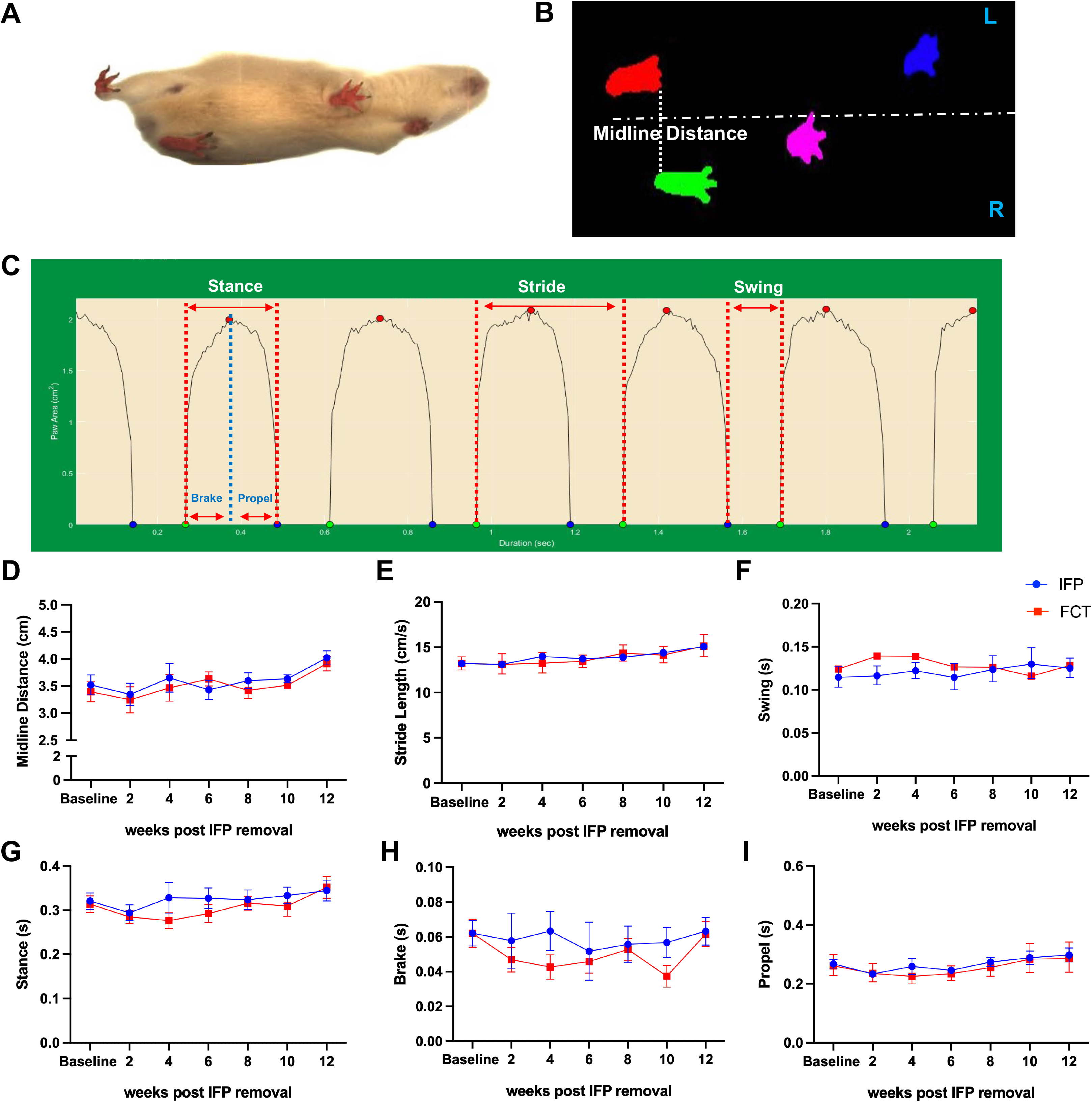
Treadmill-based gait analysis – schematics and data. [A] Video still-image showing a guinea pig running on a transparent treadmill (viewed from below) at 35 cm/s. [B] Placement for each paw is detected/illustrated from the photograph [A]. [C] Paw area in contact with the treadmill surface over time for a representative single paw (left hind limb), from which multiple stride indices can be obtained (labeled). Longitudinal data of IFP (blue, n=6) and FCT (red, n=6) limbs for midline distance [D], stride length [E], swing [F], stance [G], brake [H] and propel [I] time. P-values were determined by paired t-tests.

### Morphologic Description of H&E and Masson Trichrome Stained Slides

Haematoxylin and eosin (H&E) staining confirmed that the left (surgery sham control) knees retained the typical histological properties of IFP, including mature adipocytes, a stromal vascular fraction, and white blood cells (predominantly large and small mononuclear cells consistent with macrophages and lymphocytes, respectively) (Figure 2A). In contrast, right (IFP removal) hindlimbs exhibited development of a thick band of dense fibrous connective tissue (FCT) in the space previously occupied by the native IFP (Figure 2B). Further histological examination with Masson Trichrome stain confirmed the increased collagenous nature of the FCT compared to the native IFP (Figure 2C & D; Supplemental Figure S2).

**Figure 2.**
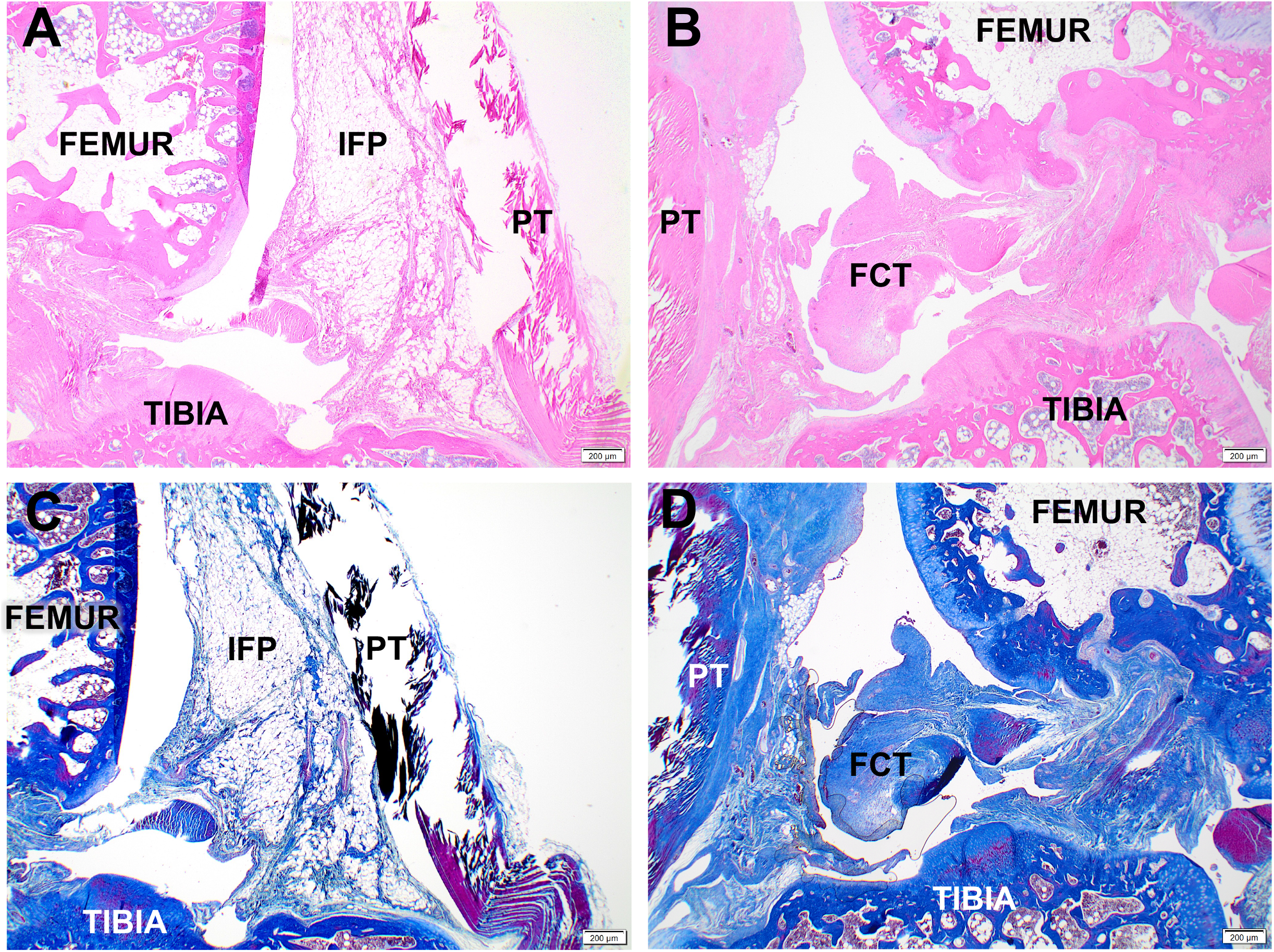
Representative mid-sagittal photomicrographs of a stifle joint from a control with the native IFP [A, C] and replacement tissue [B, D]; H&E [A, B] and Masson’s trichrome [C, D], 2x objective. [A, C] Control knee joint from a 7-month-old guinea pig depicting the normal histoanatomic location of the unaltered IFP. [B, D] Knee joint from the same 7-month-old guinea pig four months after IFP removal. The IFP space is replaced with dense fibrous connective tissue (FCT).

### MicroCT whole joint OA scores

Whole joint microCT OA scores provide a comprehensive assessment of bony changes observed in the tibia, femur, and patella of each animal. Four of the 10 IFP knees demonstrated bone sclerosis, as shown in Figure 3A (Supplemental Figure S3(C)). In addition, all IFP hind limbs presented with a mixture of small (< 1mm) and/or large (≥ 1mm) osteophytes on multiple anatomical locations (medial and/or lateral tibia, patella, or femur), with 9 out of 10 of these knees having osteophytes in two or more locations Figure 3B (Supplemental Figure S3(A)-(B)). For FCT knees, no bone sclerosis was noted. Only 3 out of 10 animals had small osteophytes present; of these 3 animals, 2 animals had small osteophytes on either the patella or femur, with 1 animal having osteophytes on both the tibia and femur Figure 3D (Supplemental Figure S3(A)-(B)). Of note, no evidence of articular bone lysis or mineralized intra-articular soft tissue were present within any knee joints. Cumulatively, OA scores were significantly higher in knees that contained the native IFP (range of 5 to 10) versus those with the replacement FCT (range of 0 to 5; *P* = 0.0020) (Figure 3E).

**Figure 3.**
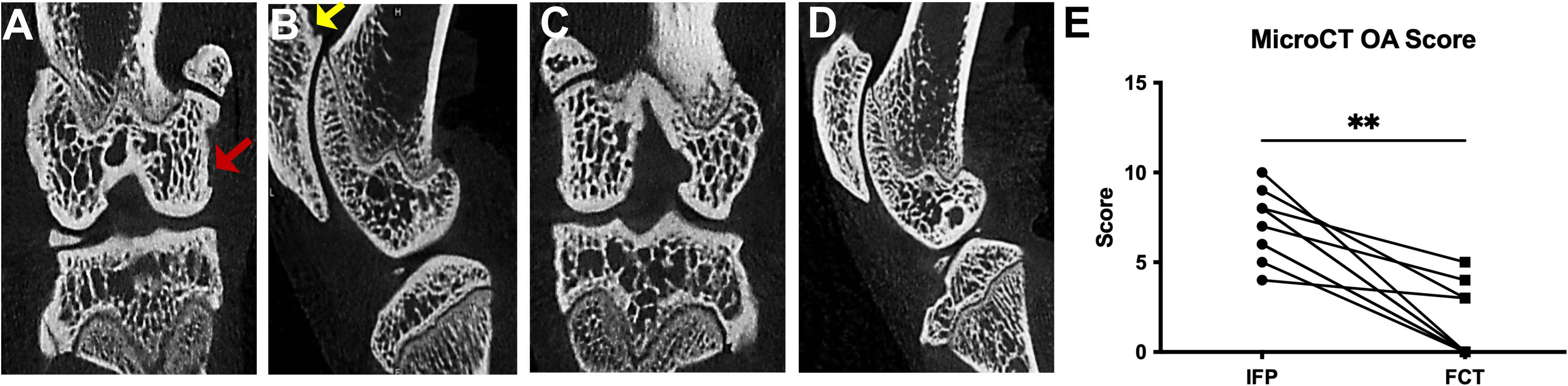
Representative photomicrographs from microCT evaluation of knee joints from the same animal using the published scoring system (Radakovich et al., *Connect Tissue Res*. 2017). [A] Coronal and [B] sagittal section of a knee containing the native IFP. Sclerosis and osteophytes are present on the medial femoral condyle (coronal section, red arrow). An osteophyte is highlighted on the proximal patella (sagittal section, yellow arrow). [C] Coronal and [D] sagittal section of the knee containing the FCT. [E] MicroCT OA score(n=10) demonstrated a significant decrease in boney changes in the FCT knees. P-value was determined by Wilcoxon matched-pairs signed rank test. *\*\*P*= *0.0020*.

### OARSI histology score

Representative lesions of both knees from the same animal are shown in Figure 4 (A) and (B). Histologic OA scores were significantly higher in IFP group compared to FCT containing knees [Figure 4 (C); *P*=0.004]. When medial and lateral compartments were analyzed independently, these compartments showed the same pattern (Supplemental Figure S4(A)-(B)). Figure 4 (A) shows irregular, undulated cartilage surface, and mild fibrillation and proteoglycan loss in the superficial zone of the tibia from the knee with the native IFP. Figure 4 (B) demonstrates a smooth cartilage surface and very mild proteoglycan loss in the knee with the FCT. Overall, the lower histological OA scores for the hind limbs that underwent IFP removal confirmed a maintenance of articular cartilage surface, proteoglycan content, and chondrocyte cellularity (Supplemental Figure S4(C)-(F)).

**Figure 4.**
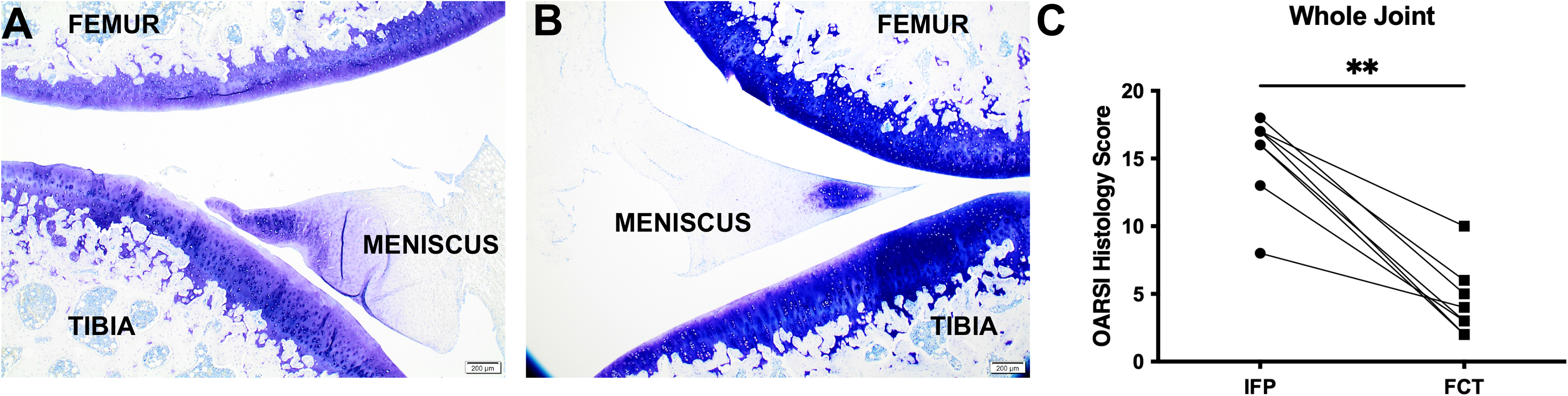
Representative photomicrographs of toluidine blue-stained sections from medial compartments for OARSI scoring. [A] Irregular articular surface with mild fibrillation and proteoglycan loss is present in the superficial zone of the tibia from the knee containing the native IFP. [B] The contralateral knee from the same animal, which contains FCT, exhibits a smooth cartilage surface and very mild proteoglycan loss. [C] OARSI whole joint score (n=8) confirmed a significant statistical difference in cartilage and proteoglycan change. P-value was determined by Wilcoxon matched-pairs signed rank test^◊^. *\*\*P*= *0.0078*

### Transcript Expression analyses

#### Components of adipose tissue

As was expected given the histologic findings, mRNA levels revealed that the FCT had a lower expression of transcripts for ADIPOQ (P = 0.0039), LEP (P = 0.0005), and FASN (P= 0.0130) than the native IFP (Figure 5 A-C).

**Figure 5.**
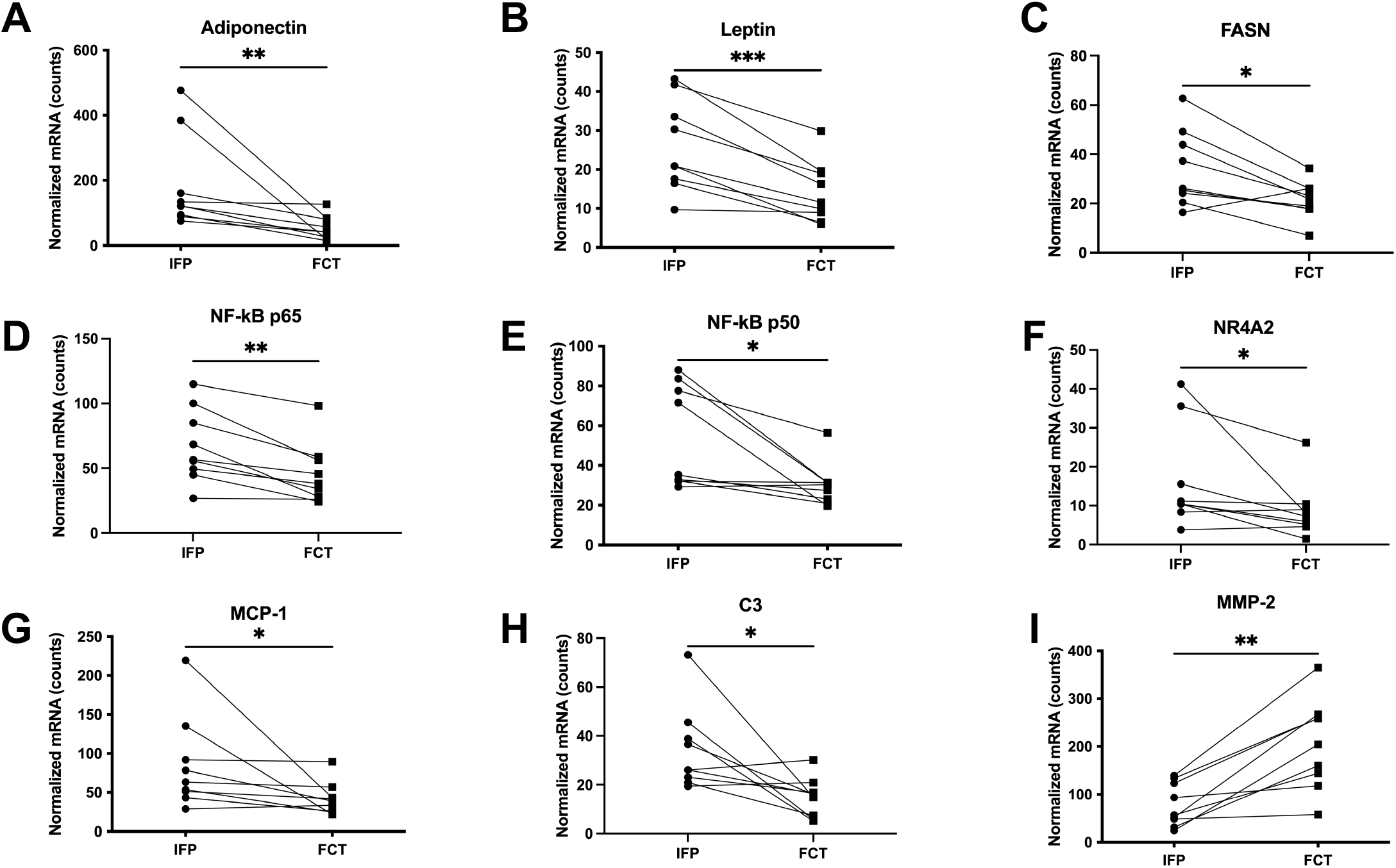
Normalized mRNA counts for adiponectin^◊^ [A], leptin^†^ [B], FASN^†^ [C], NF-kB p65^◊^ [D], NF-kBp50^◊^ [E], NR4A2^◊^ [F], MCP-1^◊^ [G], C3^◊^ [H], and MMP-2^†^ [I] in IFP versus FCT (n=9). P-values were determined by paired t-tests^†^ or Wilcoxon matched-pairs signed rank test^◊^. *\*\*\*P <0.0005, **P<0.005 and *P<0.05*

#### Inflammatory/degradative mediators

Compared to the IFP, the FCT also had decreased expression of mRNA for NF-kBp65 (P= 0.0039), NF-kB p50 (P=0.0117), NR4A2 (P=0.0273), C3 (P=0.0195), MCP-1 (P=0.0117); MMP-2 was increased (P=0.0018) (Figure 5D-I).

### IHC for Local Inflammatory Responses

To confirm the transcript expression data, MCP-1 and NF-kB p65 at the protein level were evaluated to assess local inflammation in the native IFP versus the FCT (Figures 6 & 7). Similar to the above, immunostaining of both proteins was significantly lower in the FCT (*P*<0.0001).

**Figure 6.**
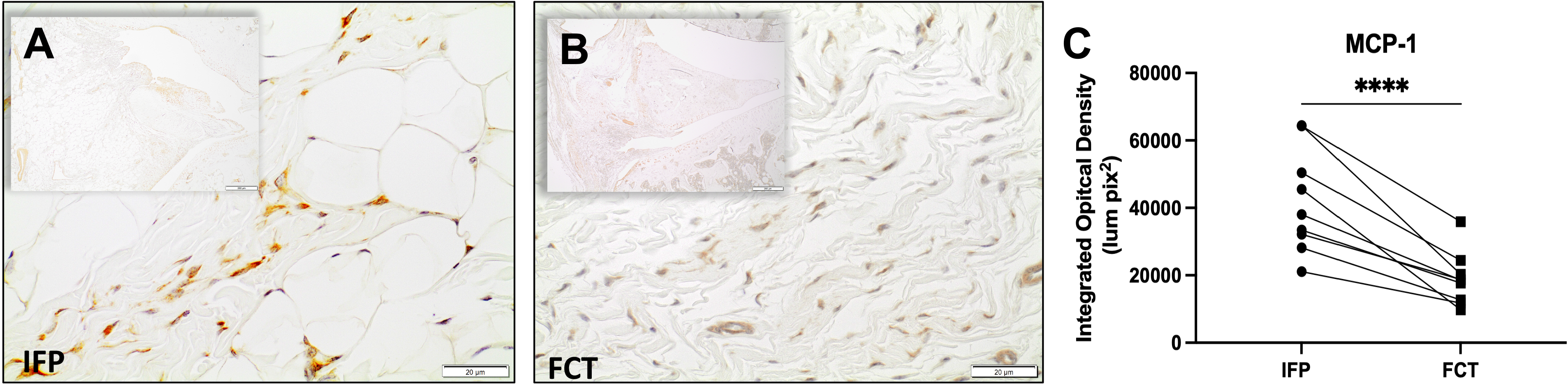
Immunohistochemistry for MCP-1 expression in [A] IFP and [B] FCT knees from the same animal (n=9); 40X objective for main photo; 20X objective for inset. [C] Removal of the IFP resulted in a decrease in MCP-1 expression relative to the FCT. Data is based on quantitative analysis of MCP-1-stained tissue subtracted from IgG control tissue for all samples. P-value was determined by paired t-test^†^. *\*\*\*\*P < 0.0001*

**Figure 7.**
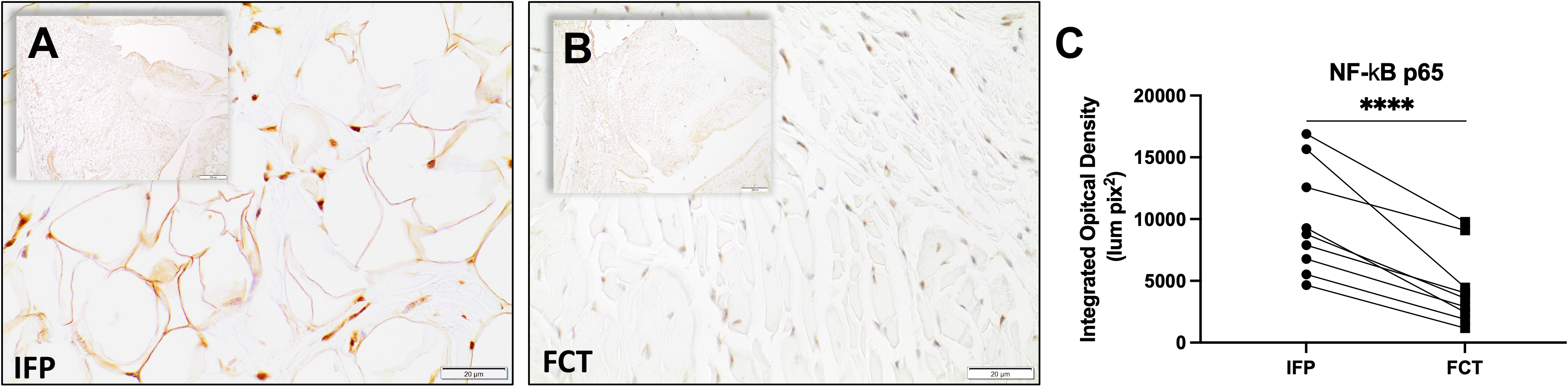
Immunohistochemistry for NF-kB p65 transcription factor protein expression in [A] IFP and [B] FCT knees from the same animal (n=9); 40X objective for main photo; 20X objective for inset. NF-kB is a transcription factor that is known for regulating inflammatory responses within inflammatory cells. [C] Compared to the native IFP, there was a decrease in p65 expression in the FCT. Data is based on quantitative analysis of p65 stained tissue subtracted from IgG control tissue for all samples. P-value was determined by paired t-test^†^. *\*\*\*\*P < 0.0001*

## Discussion

In this animal model, histomorphologic examination of the knee joints revealed that removal of the IFP resulted in the development of a thick band of collagenous FCT in the space previously occupied by the native tissue. These microscopic findings were supported by transcript expression data, which demonstrated significantly decreased expression of key adipocyte-related molecules, ADIPOQ, LEP, and FASN. Humans undergoing total knee arthroplasty combined with IFP removal have demonstrated a similar expansion/proliferation of residual tissue in the remaining void, with evidence of tissue remodeling characterized by loss of fat cells and deposition of large quantities of densely packed collagen fibers^27^. These findings are also consistent with those of *Kumer et al*.^28^, who accessed clinical and functional outcomes of Hoffa’s disease treated with high-portal arthroscopic resection and found that adipose tissue was replaced by fibrous tissue in chronic cases^28^. Additionally, these findings date back to the hallmark study conducted by Drs. Hoffa and Becker, which described this replacement process as hyperplasia and granulation of the remaining tissue such that it became interspersed with strong fibrous strings. Ultimately, endothelial cells were joined to fibrous tissue without any intervening fat^29^.

Interestingly, the current study demonstrated that early removal of the IFP prior to disease onset in an animal model of primary OA appears to have short-term benefits. Specifically, clinical microCT and histological OA scores were worse in the knee containing native adipose tissue relative to the knee with the FCT. To the authors’ knowledge, previous research in animal models of spontaneous OA have not reported the influence of IFP removal on disease pathogenesis. *Collins et al*.^30^, however, utilized a murine model of lipodystrophy that examined the direct contribution of adipose tissue in an injury model of post-traumatic osteoarthritis (PTOA) induced by destabilization of the medial meniscus (DMM). Their findings indicated that adipose tissue itself may promote PTOA susceptibility directly through adipokine signaling, triggering systemic inflammation that localizes within the joint. However, the direct mechanism in the case of PTOA remains to be determined^31^. While not analogous to our current work (as these studies focused on removal of the IFP at end stage OA), it should also be mentioned that numerous studies have investigated the effects of IFP excision in total knee arthroplasty/replacement (TKA/TKR) as it relates to Knee Society Score^32–35^. This scoring system is composed of five components: patient demographics, objective knee score (completed by the surgeon), patient expectations, patient satisfaction score, and, lastly, functional knee score (completed by the patient)^35^. To date, these studies have reported inconclusive/discrepant results in regards to either the benefit or drawback of IFP removal; recent reviews of the effect of IFP excision in TKA found no difference in anterior knee pain, range of motion, or function in the patient with OA^33–35^. Thus, our project is the first study to focus on the potential benefit of IFP removal on OA pathogenesis as a preventive therapy. In this regard, it is necessary to consider both the inflammatory and/or biomechanical benefit of the FCT versus the native IFP.

In terms of the inflammatory contribution of the IFP to OA, *Clockaerts et al*.^7^ and others^36,37^ have reported that this adipose depot, particularly in cases of obesity, can cumulatively secrete cytokines, interferons, adipokines, and growth factors, all of which exert local signaling effects on articular cartilage and synovial cells^7,36,37^. Of interest, studies have reported that individual cellular components of the IFP may contribute to OA. First, this depot serves as both a site of inflammatory/immune cell infiltration, which can provide an origin of pro-inflammatory cytokines and MMP expression. These migrating cells, including macrophages and lymphocytes, also interact and influence resident adipocytes. Second, adipocytes, themselves, are capable of secreting certain molecular markers and products able to initiate a local inflammatory response^36,37^.

Evidence to support the different functional characteristics and/or inflammatory nature of the IFP versus FCT in the current work was provided by transcript expression data and confirmatory IHC findings. As mentioned above, it is established that elevated pro-inflammatory cytokines in OA joints play a role in cartilage homeostasis^38–41^. In particular, studies have shown that NF-kB participates in many OA-associated changes, including chondrocyte catabolism, chondrocyte survival, and synovial inflammation^42,43^. Specifically, NF-kB signaling pathways mediate critical events in the inflammatory response by promoting transcription of genes encoding for cytokines and stimulating production of MMPs by synoviocytes, macrophages cells, or chondrocytes^5,42,43^. Notably, injury-induced cartilage lesions were alleviated by the knockdown of this mediator by specific small interfering RNA in animal models^44,45^. Here, we demonstrated that the FCT had downregulated NF-kB and its related nuclear orphan receptor, NR4A2, as well as NF-kB regulated genes, MCP-1 and C3. Protein expression further confirmed local decrease of one key subunit of NF-kB and MCP-1 expression within the FCT. We postulate a reduction of these inflammatory mediators may have led to decreased OA changes. Of note, MMP-2 (gelatinase A, type IV collagenase) is one of the major extracellular matrix degrading proteases and has been shown to breakdown basement membrane, which consists mainly of collagen type IV, laminin and proteoglycans^46,47^. In the current study, we found it interesting that MMP-2 expression was increased in the FCT. This finding may reflect an inherent difference in the FCT versus native adipose tissue, or may indicate that this replacement tissue was still under remodeling at the time point investigated.

It is also plausible that the FCT provided a biomechanical advantage over the IFP. Evidence for laxity of the anterior cruciate ligament has been shown in Hartley guinea pigs compared to one control strain^48^. Given this, it may also be possible that the IFP experiences similar changes in mechanical properties over time, thereby contributing to overall joint laxity and subsequent OA. As conjecture, the FCT that developed in place of the IFP may have offered improved biomechanical properties such that it protected other joint tissues. Mechanical testing on whole joints and isolated IFP or FCT may provide insight into this possibility.

A computer-aided, treadmill-based system was utilized to identify potential alterations in gait parameters in this unilateral intervention. Here, we demonstrated that IFP removal in OA-prone guinea pigs did not result in gait changes between hindlimbs across the course of this study. Previous studies have concluded that, relative to a non-OA prone strain, aged Hartley guinea pigs had shorter stride lengths and slower swing speeds^49^. In spite of our documented decrease in structural OA, removal of the IFP compared to the limb with the native IFP did not provide any changes in midline distance, stride length, swing, stance, brake or propel time. This is perhaps unsurprising, as structural joint changes are not necessarily accompanied by direct correlations to presenting clinical signs, particularly in the short-term^50^.

Considerations that should be noted in regards to this study include the fact that the current assessment involves only male animals; continued work will address findings in female guinea pigs. Further, removal of the IFP occurred prior to OA onset, which may have limited clinical application for the typical patient with OA. Indeed, examining effects of IFP removal at additional time points in the course of knee OA progression is needed to dissect the long-term benefit of IFP removal on knee health. Further, it is not known whether changes in transcript and protein expression will correlate to clinical improvement, nor if the anti-inflammatory benefits and decreased OA outcomes seen in the present work will hold true in a longer-term scenario. Finally, we acknowledge that our study design utilized the contralateral knee as a sham control to focus on within animal contrasts and comparison. It may be important to examine bi-lateral IFP removal and/or a contralateral naïve control limb to account for compensatory limb considerations. In spite of this, we have achieved a noteworthy delay of OA onset and/or progression in the Hartley guinea pig model.

## Acknowledgements

We would like to thank Crystal Richt and the staff at the University of Arizona Genetics Core.

Additionally, we would like to acknowledge the Department of Laboratory Animal Resources at Colorado State University for their compassion and commitment to these animals. The Animal Imaging Shared Resource of the University of Colorado Cancer Center receives support from P30CA046934.

## Author contributions

Maryam Afzali, Lauren Radakovich, and Kelly Santangelo take responsibility for the integrity of the work.

Study conception and design: KSS, LBR

Collection and assembly of data: MFA, LBR, MAC, JLS, MMS, AJM

Analysis and interpretation of the data: MFA, LBR, MAC, JLS, MMS, THD, KSS

Drafting of article: MFA, LBR

Critical revision of the article for important intellectual content: LBR, MAC, JLS, MMS, AJM, THD, KSS

Final approval of the article: MFA, LBR, MAC, JLS, MMS, AJM, THD, KSS

Obtaining of funding: KSS

## Supplemental Material

**Supplemental Figure S1.**
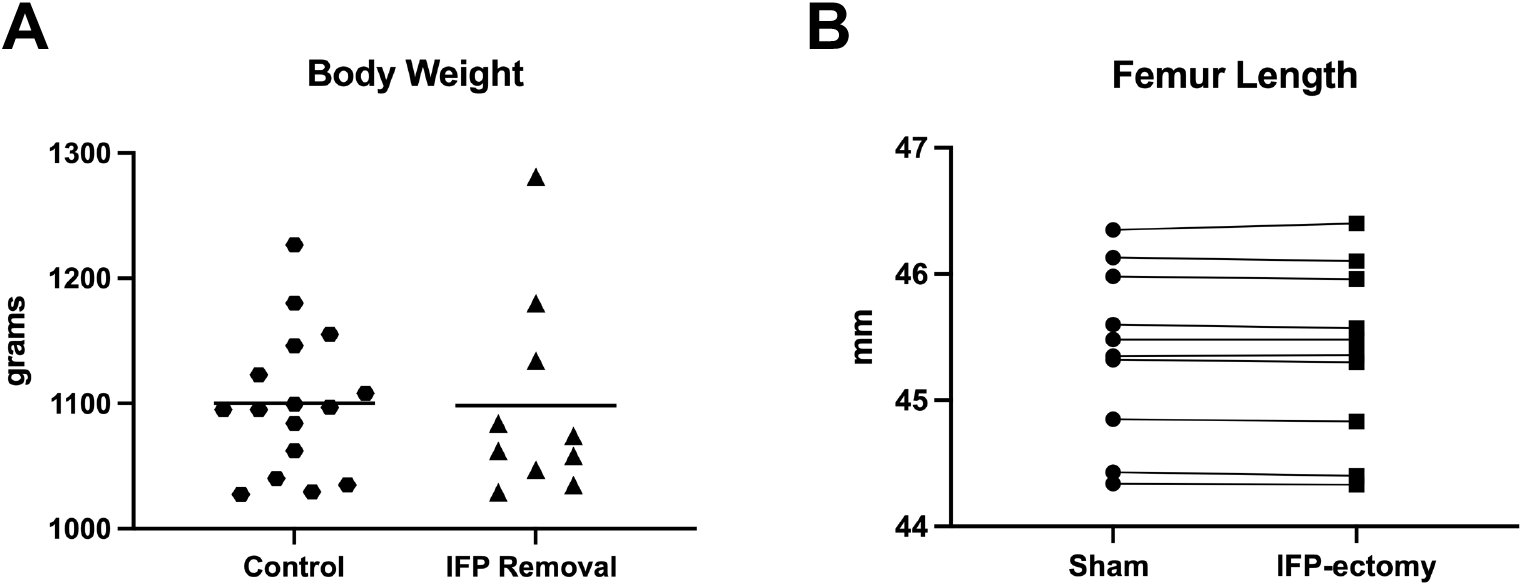
Body weight [A] in 7-month Hartley guinea pig controls (n=16) and IFP removal groups (n=10). [B] Femur lengths (n=10) between IFP and FCT-containing limbs. Black line represents mean value.

**Supplemental Figure S2.**
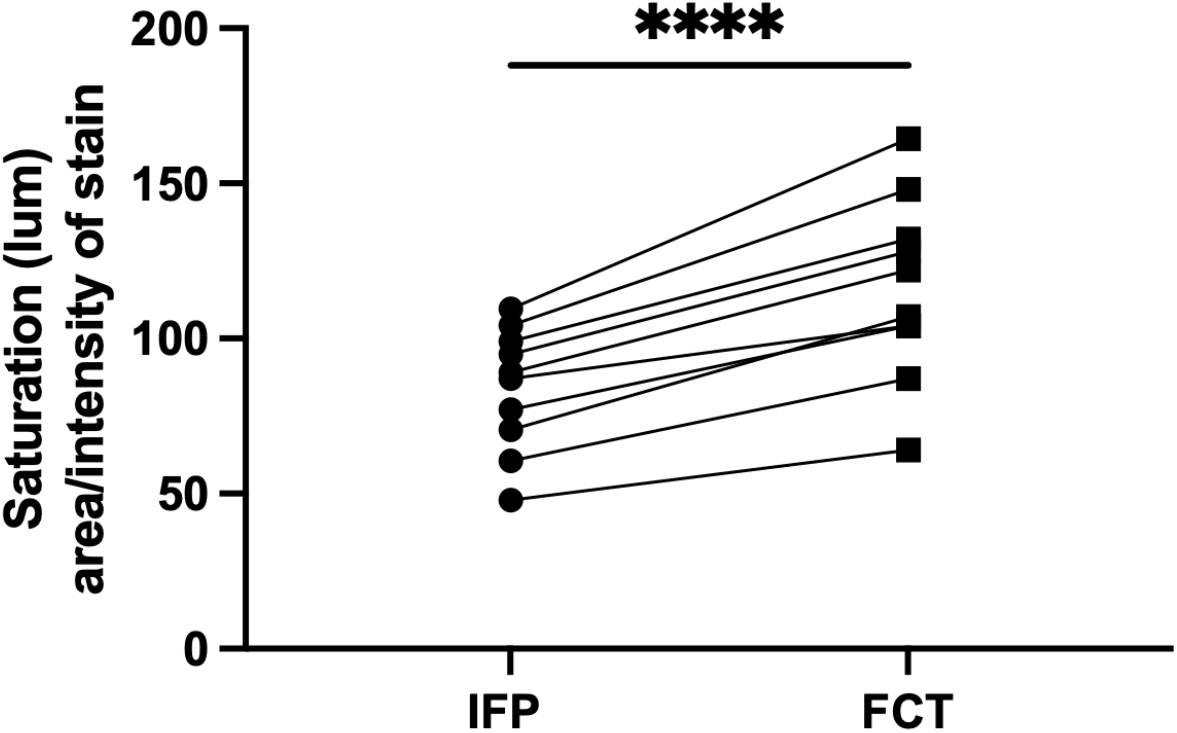
Quantitative analysis of Masson’s Trichrome staining in IFP vs FCT (n=10). P-value was determined by paired t-test^†^. *\*\*\*\*P < 0.0001*

**Supplemental Figure S3.**
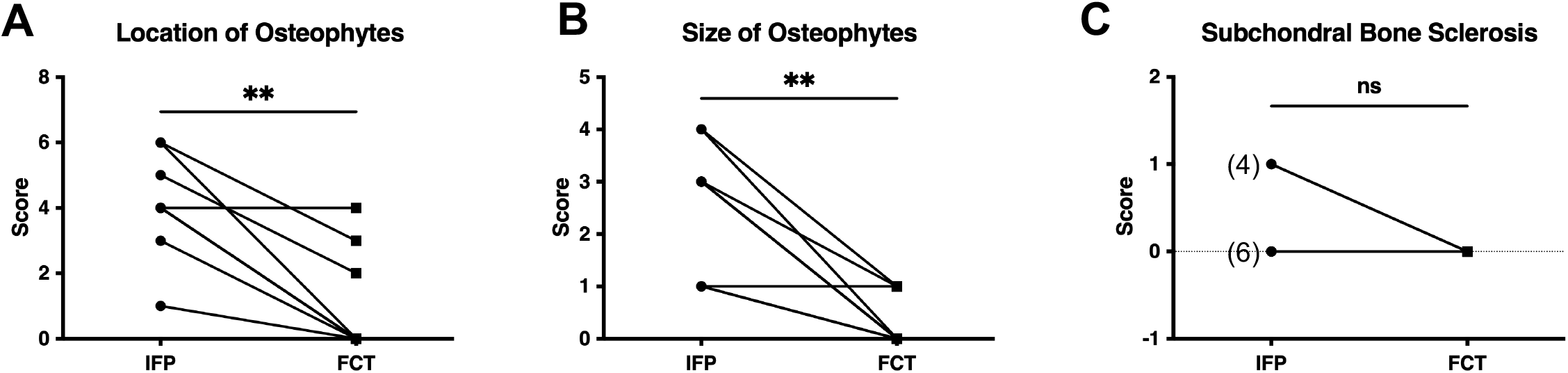
Contributions to whole joint MicroCT OA score components. [A] Location of osteophytes^◊^ (n=10, possible range of scores 0–9). Mean location of osteophytes score in the FCT was 0.900 (n = 10; 95% CI −0.1901-1.990) and 4.100 in the IFP (*n* = 10; 95% CI 3.063–5.137). [B] Size of osteophytes^◊^ (n=10; possible range of scores 0-4). Mean size of osteophytes score in the FCT was 0.300 (n= 10; 95% CI −0.046–0.645) and 2.60 in the IFP (n =10; 95% CI 1.760–3.440). [C] Subchondral bone sclerosis^◊^ (n=10; possible score of 0 for absence, 1 for presence). Mean score for subchondral bone sclerosis was 0.00 in the FCT (n=10; 95% CI 0 – 0) and 0.4 in the IFP (n = 10; 95% CI 0.031–0.764). Data with non-Gaussian distribution were compared using non-parametric Wilcoxon matched – pairs signed rank test^◊^. *\*\*P=0.0039*

Supplemental Figure S4. Contributions of individual components to whole joint OARSI score. [A] Medial compartment^◊^ OARSI score (possible range of scores 0–42). Mean OARSI histologic score in the medial compartment of the FCT was 2.63 (n = 8; 95% CI 1.08-4.17) and 11.63 in the IFP (n = 8; 95% CI 8.28–14.97); *\*\*P*=*0.078*. [B] Lateral compartment^◊^ OARSI score (possible range of scores 0–42). Mean OARSI histologic score in the lateral compartment of the knee was 3.63 in the IFP (n= 8; 95% CI 2.29–4.96) and 1.75 in the FCT (n=8; 95% CI 0.68–2.82). [C] Articular cartilage structure (n=8; possible range of scores 0–32 for the sum of all compartments). Mean score for articular cartilage structure was 4.38 in the IFP (*n* = 8; 95% CI 2.77 – 5.98) and 1.63 in the FCT (n = 8; 95% CI 0.86–2.39); *\*P*=*0.0293*. [D] Proteoglycan content^†^ (n=8; possible range of scores 0–24 for the sum of all compartments). Mean score for proteoglycan content was 7.00 in the IFP (n = 8; 95% CI 5.66–8.34) and 2.50 in the FCT (n = 8; 95% CI 0.45–4.55); *\*\*P*=*0.0016*. [E] Cellularity^†^ (possible range of scores 0–12 for the sum of all compartments). Mean score for cellularity was 4.13 in the IFP (*n* = 8; 95% CI 2.68-5.57) and 0.88 in the FCT (n = 8; 95% CI −0.07 – 1.82); *\*\*P*=*0.0015*. [F] Tidemark integrity^◊^ (possible range of scores 0–4 for the sum of all compartments). Mean score for tidemark integrity was 0.25 in the IFP (n = 8) and 0 in the FCT (n = 8). Two animals were unable to be evaluated for whole joint OARSI histologic grading due to appropriate tissues sections being unavailable. Normally distributed data with similar variance were compared using paired t-tests^†^. Data with non-Gaussian distribution were compared using Wilcoxon matched-pairs signed rank test^◊^.Click here to access/download

**Supplemental Table 1.**
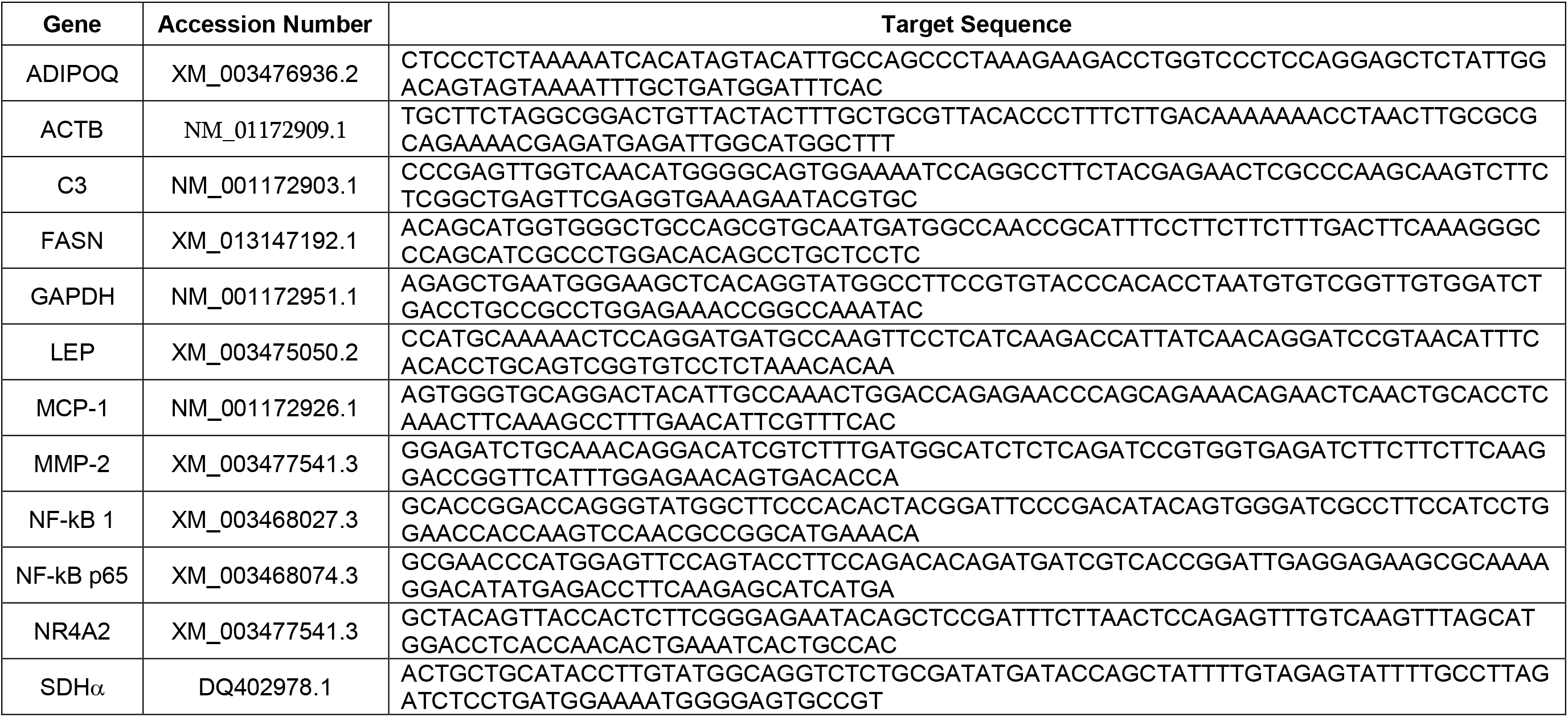
Primer sequences used for Nanostring gene expression analysis. The length of all target sequences are 99 base pairs.

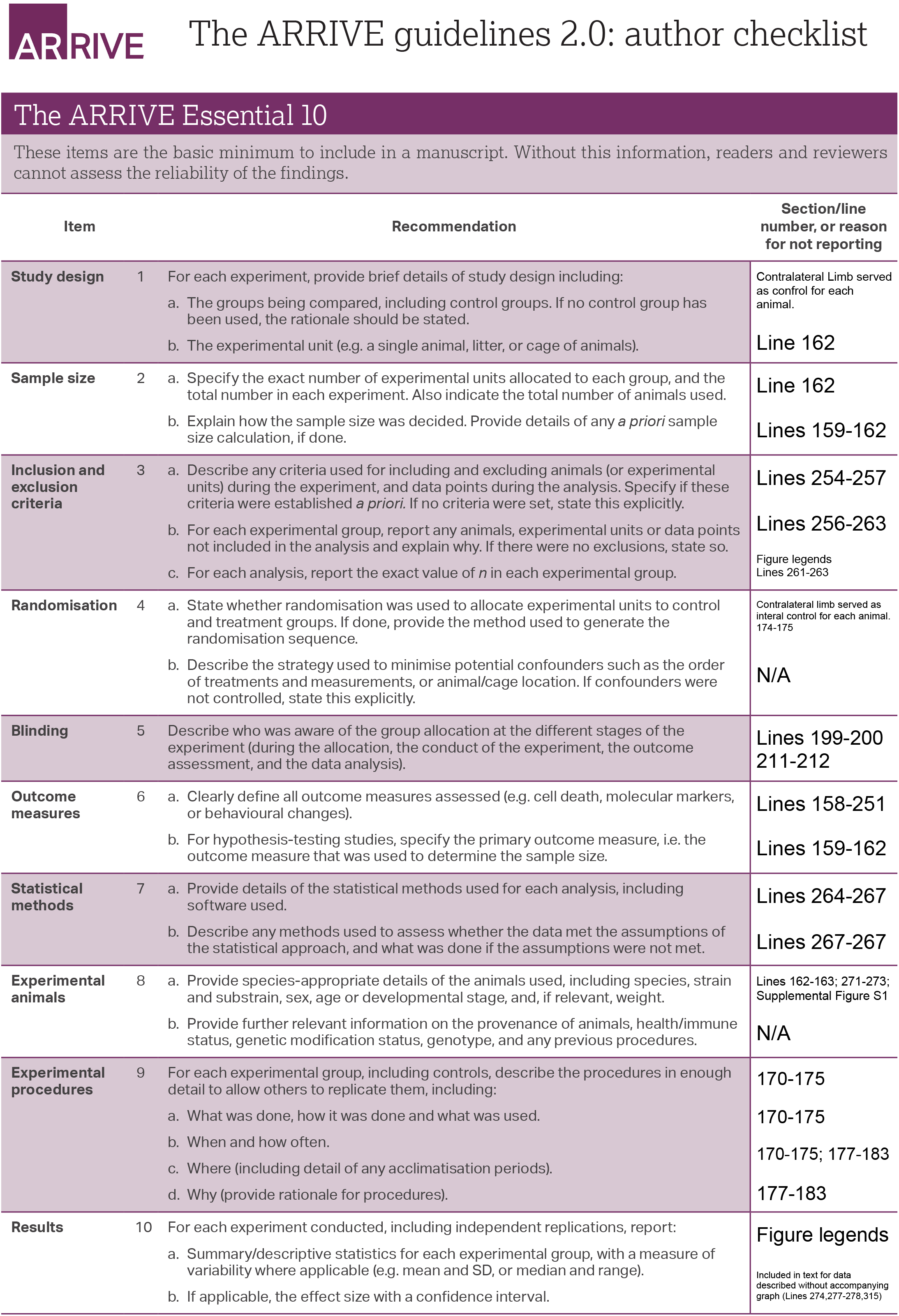

